# Metabolic-sensing in AgRP neurons integrates homeostatic state with dopamine signalling in the striatum

**DOI:** 10.1101/2021.03.22.436393

**Authors:** Alex Reichenbach, Rachel E Clarke, Romana Stark, Sarah Haas Lockie, Mathieu Mequinion, Felicia Reed, Sasha Rawlinson, Harry Dempsey, Tara Sepehrizadeh, Michael DeVeer, Astrid C Munder, Juan Nunez-Iglesias, David C. Spanswick, Randall Mynatt, Alexxai V. Kravitz, Christopher V. Dayas, Robyn Brown, Zane B. Andrews

## Abstract

Hunger increases the motivation of an organism to seek out and consume highly palatable energy dense foods. While hunger-sensing Agouti-related peptide (AgRP) neurons influence this process, whether metabolic detection of homeostatic state via metabolic sensing in AgRP neurons potentiates motivation through the midbrain dopamine system is unexplored. Here, we used the AgRP-specific deletion of carnitine acetyltransferase (Crat), a metabolic enzyme regulating glucose and fatty acid oxidation, as a model of impaired metabolic-sensing in AgRP neurons. We then tested the hypothesis that appropriate metabolic-sensing in AgRP neurons is required to increase food reward motivation by modulating accumbal or striatal dopamine release. Electrophysiological studies confirm that Crat deletion in AgRP neurons (KO) impairs normal *ex vivo* glucose-sensing, and *in vivo* photometry experiments show that AgRP neurons in KO mice do not exhibit normal responses to repeated palatable food presentation and consumption, highlighting that this model is appropriate to test the hypothesis. Fiber photometry experiments, using the dopamine sensor GRAB-DA, revealed that impaired metabolic-sensing reduces acute dopamine release (seconds) in the nucleus accumbens, but not the dorsal striatum, to palatable food consumption and during operant responding. Positron electron tomography (PET) methods indicated that impaired metabolic-sensing in AgRP neurons suppressed radiolabelled 18F-fDOPA accumulation after ∼30 minutes in the dorsal striatum but not the ventral striatum, suggesting a role for AgRP neurons to restrict a long term post-ingestive dopamine response in the dorsal striatum. Finally, impaired metabolic-sensing in AgRP neurons suppresses motivated operant responding for sucrose rewards. Notably, these behavioural effects are potentiated in the hungry state and therefore highlight that metabolic-sensing in AgRP neurons is required for the appropriate temporal integration and transmission of homeostatic hunger-sensing to dopamine signalling in the striatum.

## INTRODUCTION

The motivation to approach and consume food depends not only on the palatability and caloric density of the available food source, but also on the energy state of the organism. For example, when food is abundant many prey species forage within known territories to reduce survival threats (*1*). Conversely, when food is scarce, animals are motivated to take greater risks and forage within unfamiliar territories to search for food (*2*). Thus, the motivation to seek palatable, energy-dense food evolved as a key mechanism for survival and maturation in an environment with limited food availability. Given that heightened motivation for palatable food in an environment of low food availability shaped an evolutionary benefit, it is not surprising that homeostatic feeding circuits can have a profound effects on motivation. In today’s society with high caloric foods readily available, these circuits may contribute to the overconsumption of highly palatable and calorie dense foods, which is a leading cause for today’s obesity crisis. Indeed, human evidence shows that fasting biases reward systems to high caloric foods (*3, 4*).

Agouti-related peptide (AgRP) neurons in the arcuate nucleus of the hypothalamus (ARC) are a critical population of hunger-sensitive neurons that primarily function to increase appetite or conserve energy (*5–9*). A key element of appetitive behavior is the increased motivation for goal-directed outcomes. Intriguingly, hunger has been used for decades in behavioral neuroscience to improve performance and learning in operant tasks. AgRP neurons function within this framework by increasing the willingness to work for food and food rewards to the same level as that seen in fasted mice (*7, 10–12*). Moreover, AgRP neurons influence motivation circuits (*13–15*), with chemogenetic activation elevating dopamine release in response to food consumption (*13*). Given that AgRP neurons are most active during periods of energy deficit (*16–20*), we hypothesized that they transmit hunger-specific metabolic information to motivation circuits. However, whether AgRP neurons are required to gauge homeostatic state though metabolic-sensing, in order to influence motivation neural circuits, remains unknown.

Sensory cues that predict food availability and palatability rapidly suppress hunger-sensitive AgRP neuronal activity in hungry mice (*11, 18, 21, 22*). Although this decrease in activity occurs prior to food consumption, post-ingestive caloric feedback is required to sustainably reduce AgRP neuronal activity (*22, 23*). Indeed, the reduction of AgRP neuronal activity correlates with the number of calories ingested and was not observed after repeated ingestion of a non-caloric sweetened gel (*22*). These studies highlight that the maintenance of normal AgRP activity in response to post-ingestive gastrointestinal feedback requires metabolic sensing of available calories in combination with gut-brain neural communication (*22–24*). Previously we demonstrated that carnitine acetyltransferase (Crat) in AgRP neurons was required as a molecular sensor for peripheral substrate utilisation during fasting and refeeding (*25*). Moreover, Crat in AgRP neurons programmed a broad metabolic response of the AgRP proteome and was required to promote normal refeeding after fasting (*25*). The reduced feeding response after fasting in KO mice suggested that metabolic sensing of homeostatic state by AgRP neurons transmits hunger-specific metabolic information into neural circuits controlling dopamine signalling and the motivational aspects of food-directed behaviour. Here we describe experiments demonstrating that impaired metabolic-sensing in AgRP neurons, using the conditional deletion of Crat from AgRP neurons as a validated model, reduces motivated behaviour during an operant task and dopamine release in the nucleus accumbens (NAc) or dorsal striatum, over different time frames, in response to palatable food rewards. These studies highlight that metabolic-sensing in AgRP neurons is required for the appropriate temporal integration of hunger-sensing to potentiate food reward-related dopamine release in the striatum and control motivation for palatable food rewards.

## Methods

### Animals

All experiments were conducted in compliance with the Monash University Animal Ethics Committee guidelines. Male and female mice were kept under standard laboratory conditions with free access to food (chow diet, catalog no. 8720610, Barastoc Stockfeeds, Victoria, Australia) and water at 23C in a 12-hr light/ dark cycle and were group-housed to prevent isolation stress unless otherwise stated. All mice were aged 8 weeks or older for experiments unless otherwise stated. AgRP-ires-cre mice were obtained from Jackson Laboratory AgRP^tm1(cre)Low/J^ (stock no. 012899) and bred with NPY GFP mice (B6.FVB-Tg(Npy-hr*GFP*)1Lowl/J; stock number 006417; The Jackson Laboratory, Maine, USA). AgRP-ires-cre::NPY GFP mice were then crossed with Crat^fl/fl^ mice donated by Randall Mynatt (Pennington Biomedical Research Center, LA, USA) in order to delete Crat from AgRP neurons (AgRP^cre/wt^::Crat^fl/fl^ mice; designated as KO). AgRP^wt/wt^::Crat^fl/fl^ littermate mice were used as control animals (designated as WT). For in vivo photometry AgRP^cre/wt^::Crat^wt/wt^ mice; designated as WT to allow for cre-dependent expression of GCaMP6s specifically in AgRP neurons (KO same as above).

### Electrophysiology

Animals were fasted overnight and 250 µm thick coronal hypothalamic brain slices containing the ARC were prepared from 10 male AgRP Crat KO and 9 WT mice (8-12 weeks) expressing GFP in NPY neurons, and stored at room temperature before transferral to the recording chamber. Slices were continuously superfused at 4-5 ml/min with oxygenated (95% O_2_, 5% CO_2_) artificial cerebrospinal fluid (aCSF) of the following composition (in mM): NaCl 127, KCl 1.9, KH_2_PO_4_ 1.2, CaCl_2_ 2.4, MgCl_2_ 1.3, NaHCO_3_ 26, D-Glucose 2, Mannitol 8 (310 mOsm, pH 7.4). Hypothalamic neurons were visualised under IR illumination using a 63x or 40x water immersion objective on an upright microscope (Axioskop 2, Zeiss) and an Axiocam MRm camera (Zeiss). AgRP neurons were identified using a GFP filter set. Patch pipettes (8-11 MΩ) were pulled from borosilicate glass capillaries (Harvard Apparatus) and filled with intracellular solution containing (in mM): K-gluconate 140, KCl 10, EGTA 1, HEPES 10, Na-ATP 2, Na-GTP 0.3 (300 mOsm and pH 7.3, adjusted with KOH). Whole-cell current clamp recordings were made using the MultiClamp 700B amplifier, digitized with the Digidata 1550B interface, and acquired in pClamp 10.6 at 5 kHz sampling rate (Axon Instruments). To test the influence of an elevated extracellular glucose concentration on neural activity, aCSF was prepared as described above with the following changes: D-Glucose 5mM, Mannitol 5mM, and bath-applied for 10-15min. Data were analysed in Clampfit 10.6 (Axon Instruments) and plotted in Graphpad Prism 8.3. Figures were further prepared in Adobe Illustrator CC 2020.

### Fiber photometry

Mice for fiber photometry experiments were anesthetised (2-3% isoflurane) and injected with Metacam (5 mg/kg) prior to placing into stereotaxic frame (Stoelting/Kopf) on heatpad (37C) and viral injections were performed as previously described(*26*). Cre-dependent GCaMP6s (Addgene #100845 AAV9-hSyn-FLEX-GCaMP6s-WPRE-SV40) was unilaterally injected into the ARC (−1.5mm Bregma; +/- 0.2 mm lateral; −5.6mm ventral from surface of brain). Non-cre dependent dopamine sensor (YL10012-AAV9: AAV-hSyn-DA4.3)(*27*) was unilaterally injected in dorsal striatum (bregma 0.5mm, midline 1.3 mm. skull −3.4mm) or NAc (bregma 1.2mm, midline 0.5 mm. skull −4.8mm). Injections were 150nl/side @25nl/min, 5 min rest and ferrule capped fibers (400µm core, NA 0.48 Doric, MF1.25 400/430-0.48) implanted above injection site and secured with dental cement (GBond, Japan). Mice had 2 weeks recovery before commencement of experiments.

All fiber photometric recordings were performed with a rig using optical components from Doric lenses controlled by Tucker Davis Technologies fiber photometry processor RZ5P. TDT Synapse software was used for data acquisition. Prior to experiments, baseline GCaMP6s or dopamine signal was measured and LED power was adjusted for each mouse to achieve approximately 200mV for 465nm (530 Hz) and 100mV for 405nm (210 Hz). 405nm was used as an isosbestic control, which is wavelength at which GCaMP6 excitation is independent from intracellular [Ca^2+^], allowing for an assessment of motion artefact. For data analysis we used a modified python code provided by TDT(*28, 29*). The modified code is available at github/AlexReichenbach upon publication. Briefly, raw traces were down sampled and df/f (f465nm-f405nm/f405nm) was calculated to detrend signal bleaching and remove any motion artefact. Individual z-scores around each precisely timed transistor-transistor logic (TTL)-triggered event were extracted to allow a standardised comparison between events. Z-score normalisation indicates the number of standard deviations a particular data point is away from a defined baseline mean and was calculated according to the following equation (s. = [‘:f – ‘:f^]/’2}; where the raw data point ‘:f is subtracted from the mean of the baseline period ‘:f^ divided by the standard deviation ‘2 of the baseline period. The baseline period was defined as the period prior to a defined TTL-triggered behavioural event as described in results and figure legends.

To measure dopamine release to non-food-object/chow/peanut butter chip, single-housed mice with fiber implants were habituated to receiving Reese’s® peanut butter chips in the home cage (Macronutrient composition – Fat 29%, Carbohydrate 52%, Protein 3%). On the test day mice were connected to fiber photometry setup in their home cage and a small ceramic bowl placed inside. Recording started after 5 minutes acclimation period. In 2 minute intervals a small wood dowel (novel non-food object), a chow pellet, and a peanut butter chip were dropped into the ceramic bowel in that order. Mice were fed or fasted in random cross over design. For fiber photometry using GCaMP6s in AgRP neurons, successful targeting of AgRP neurons was tested 2 weeks after surgery by injecting ghrelin (1 mg/kg). Baseline GCaMP6s signals were recorded 15 min prior to ip injection and mice presented with standard chow 90 min after injection. To measure AgRP responsiveness to palatable food, naïve mice were presented with Reese’s® peanut butter chips. Multiple trails of this experiment were repeated over multiple days in fed and fasted mice (overnight fasting ∼16 hours).

### Operant conditioning task

For operant conditioning experiments, Feeding Experiment Devices version 3 (FED3)(*30*) were placed overnight (16h) inside home cages under *ad libitum* conditions trained to reliably nose poke on fixed ratio (FR)1, FR3 and FR5 schedules (criteria to move to higher schedule was 3 consecutive days over 75% correct nose pokes) The dispensing of a sugar pellet (20mg, 65% sucrose, 5TUT Test Diets, Saint Louis, Missouri, USA) was paired with an LED light cue. During these FR sessions, a nose poke in the ‘active’ hole resulted in the delivery of a sugar pellet and was paired with a LED light cue whereas a nose poke in the ‘inactive’ hold resulted in no programmed response. Importantly, mice were never food restricted during training to prevent this confounding the interpretation of our results. Once stable operant responding was established (> 75% correct nose pokes on FR5) mice were placed on a progressive ratio (PR) schedule under *ad libitum* fed conditions for a single overnight PR session (16 h). PR sessions were based on a Richardson Roberts schedule(*31*) where the number of pokes required to obtain a reward increased after every completed trial in the following pattern; 1, 2, 4, 6, 9, 12, 15, 20, 25, 32, etc. After this single PR session, stable FR5 responding was re-established (>75% correct nose pokes, typically 1 FR session) and a second PR session was performed except this time mice were fasted 16h during to the PR session. Separate cohorts with fiber implants in the dorsal striatum or NAc were trained to receive sugar rewards as described above and dopamine responses to both nose pokes and sugar pellet retrieval were recorded in fed state or after an overnight fast.

### PET/CT and MRI

Mice were single housed and trained to receive a single Reese’s^®^ peanut butter chip 6 hours into the light phase (time of PET scan) for 1 week. The day before PET/CT scans, mice were fasted overnight (18 hours) with ad lib access to water. On the experimental day, mice were injected IP with 10mg/kg Benserazide (Selleckchem.com) and 10mg/kg Entacapone (Selleckchem.com) 30 minutes before injecting radiolabelled fDOPA (approx. 5MBq) into the tail vein. After 5 min rest, mice received 1 peanut butter chip and only those mice that initiated feeding within the first minute were included in the analysis. Then mice were anaesthetised (1-2.5% isoflurane) and a 65 min PET/CT scan was performed under anaesthesia. The scan was acquired using the Inveon scanner (Siemens Inveon). CT was generated using the following parameters: 97µm of resolution, 80kV voltage and 500uA current, mainly for attenuation correction and MRI overlaying purposes. PET scans were acquired for 60 minutes (16 time frames). One week later MRI scan was performed on these animals to generate T2 weighted images using these parameters: 3D Flash sequence, TE/TR=8/60ms, 4 averages, flip angle= 10 degree, resolution = 0.155 mm^3^, scan time 20 minutes. PET images were analysed using IRW software (Inveon Research Workplace 4.2). PET/CT images were overlaid manually with the T2-Weighted images. Region of Interests (ROIs) were generated carefully by one investigator (AR) on the T2 images on left/right ventral striatum, left/right dorsal striatum and cerebellum (according to Allen brain atlas) and PET voxel intensities (Unit Bq/ml) at 3 different time points were exported for further analysis. We specifically choose to use the ventral striatum, rather than the NAc, due to greater accuracy with ROI identification. SUV (Standardized Uptake Value) of ROIs were calculated for each time point using the following equation: SUV= C_PET_(T)/(Dose/Weight). C_PET_(T) = Tissue radioactivity concentration at time T. Dose= administration dose at time of injection (decay corrected to the time points). Weight= animal body weight. To eliminate bias for slight variation of size/location of ROI we used upper bound Bq/ml values to calculate SUVmax(*32*) and normalised to cerebellum as reference: SUVRmax= SUVmax target/SUVmax reference.

### Immunohistochemistry

To confirm viral injection and fiber placements, animals were deeply anesthetized with isoflurane and perfused with 0.05 M PBS, followed by 4% paraformaldehyde. Brains were postfixed in 4% paraformaldehyde overnight at 4°C then placed in 30% sucrose. Brains were cut at 40 μ on a cryostat, and every fourth section was collected and stored in cryoprotectant at −20°C. Sections were washed in 0.1 M phosphate buffer (PB), incubated with 1% hydrogen peroxide (H_2_O_2_) for 15 minutes to prevent endogenous peroxidase, activity, and blocked for 1 hour with 5% normal horse serum (NHS) in 0.3% Triton in 0.1 M PB. Sections were incubated with chicken anti-GFP (ab13970, Abcam) at 1:1000 in diluent of 5% NHS in 0.3% Triton in 0.1 M PB. After incubation, the sections were washed and incubated with Alexa Fluor goat anti-chicken 488 antibody (Invitrogen) at 1:500 in 0.3% Triton in 0.1 M PB. Sections were then washed, mounted, and coverslipped.

### Statistical analysis

Statistical analyses were performed using GraphPad Prism for MacOS X. Data are represented as mean ± SEM. Two-way ANOVAs with post hoc tests were used to determine statistical significance. A two-tailed Student’s paired or unpaired t-test (see figure legends for specific details) was used when comparing genotype only. p < 0.05 was considered statistically significant.

## RESULTS

### Crat deletion in AgRP neurons is a valid model of impaired metabolic sensing

In order to demonstrate that Crat deletion in AgRP neurons is a reliable model of impaired metabolic-sensing, we prepared hypothalamic brain slices from WT and KO mice for electrophysiological characterisation of glucose sensing in AgRP neurons. We detected no differences in the fundamental electrophysiological properties of AgRP neurons including resting membrane potential (Fig 1A), input resistance (Fig 1B), spontaneous firing frequency (Fig 1C), although peak action potential amplitude, was significantly lower in KO mice (Fig 1D). Collectively, these studies demonstrate that Crat in AgRP neurons has little effect on the intrinsic electrophysiological properties of the cell. Similarly, no genotype-dependent changes in the frequency of spontaneous excitatory or inhibitory post-synaptic potentials (EPSPs or IPSPs, respectively) were detected (Supp Fig 1A-B), indicating no differences in the external synaptic input in WT compared to KO mice. To show impaired glucose-sensing in AgRP neurons, we recorded neuronal responses to an increase in extracellular glucose from a basal level of 2mM to 5mM. These glucose concentrations were chosen to represent brain glucose concentrations under fasting and fed conditions, based on glucose concentration estimated in the CSF from fasted and fed rats and mice (*33*). Glucose-excited neurons were defined based upon a response characterised by membrane potential depolarization and/or an increase in action potential firing frequency with increased extracellular glucose (Supp Fig1C). Glucose-inhibited cells were identified by responses to increased glucose characterised by membrane potential hyperpolarisation and/or a reduction in spontaneous action potential firing frequency (Supp Fig1D). In WT mice, 48% of cells (n=10/21) were excited, 33% of cells (n=7/21) were inhibited and 19% (n=4/21) were insensitive to changes in extracellular glucose (Fig 1E). In KO mice, the number of glucose-excited cells was reduced compared to WT, with only 28% classified as glucose-excited (n=7/25). Similarly, the incidence of glucose-inhibited cells was reduced to 16% (n=4/25) of the population compared to WT mice and the majority of cells not responding to 5mM glucose 56% (n=14/25) and classified as glucose insensitive (Fig 1F). In total 17/21 AgRP neurons responded to an increase in extracellular glucose in WT mice, whereas as only 11/25 AgRP neurons responded to glucose in KO mice, representing a decrease in glucose-responsive neurons (WT 81% vs KO 44%). To examine whether these *ex vivo* recordings had an impact on AgRP neurons *in vivo,* we measured GCaMP6 fluorescent in freely moving WT and KO mice, as an index of neural activity using fiber photometry. Fed WT or KO mice were presented with a novel peanut butter chip (PB) and the fall in AgRP activity was measured, as previously described (*11, 18, 21*). The fall in AgRP activity was similar in WT and KO mice in response to first PB exposure (Fig 1G-H), and with repeated exposure across different sessions WT mice showed a greater suppression of activity relative to first exposure (Supp Fig 1E; Fig 1L), consistent with previous studies showing that the magnitude of activity suppression is proportional calories consumed (*22*). Unlike WT mice, KO mice did not exhibit any further reduction in AgRP activity to repeated PB exposure (Supp Fig 1F, Fig 1L). The reduction in KO mice to repeated PB exposure was significantly attenuated compared to WT mice at all time points after PB exposure either in ad libitum (Fig 1J-K) or fasted mice (Fig 1M-N). These results suggest KO mice could not integrate caloric information associated with PB consumption either in the fed or fasted state and therefore did not show the expected reduction in activity, which is known to be proportional to calorie content (*22*). To determine if this response was specific to metabolic sensing of calorie content and not a generalised response to due impaired AgRP neuronal function caused by Crat deletion, we examined ghrelin-induced food intake and AgRP neural activity (Supp Fig 2). IP injection of ghrelin (1 mg/kg) significantly increased food intake after 4 hours (Supp Fig 2B; main effect of ghrelin), however no differences in genotype were observed. IP ghrelin significantly increased AgRP neural activity (Supp Fig 2D-E; main effect of ghrelin) however no differences in genotype were observed and the presentation of chow diet to ghrelin-injected mice produced an equal suppression in AgRP neural activity (Supp Fig 2F-G; main effect of food access), again without differences between WT and KO mice. Finally, we observed no differences in anxiety-like behaviour in the light dark box or elevated plus maze (Supp Fig I-L). The lack of genotype differences in ghrelin-induced food intake and AgRP activity, or differences in anxiety-like behaviour, which can be modulated by AgRP neurons (*34*), suggests the deletion of Crat from AgRP neurons does not cause a global deficit in function. Taken together, the specific impairments in *ex vivo* glucose sensing and *in vivo* responding to palatable calorie consumption show that Crat deletion in AgRP is a useful model to assess the impact of impaired metabolic-sensing in AgRP neurons on motivated behaviour and dopamine release in response to food rewards.

**Figure 1:**
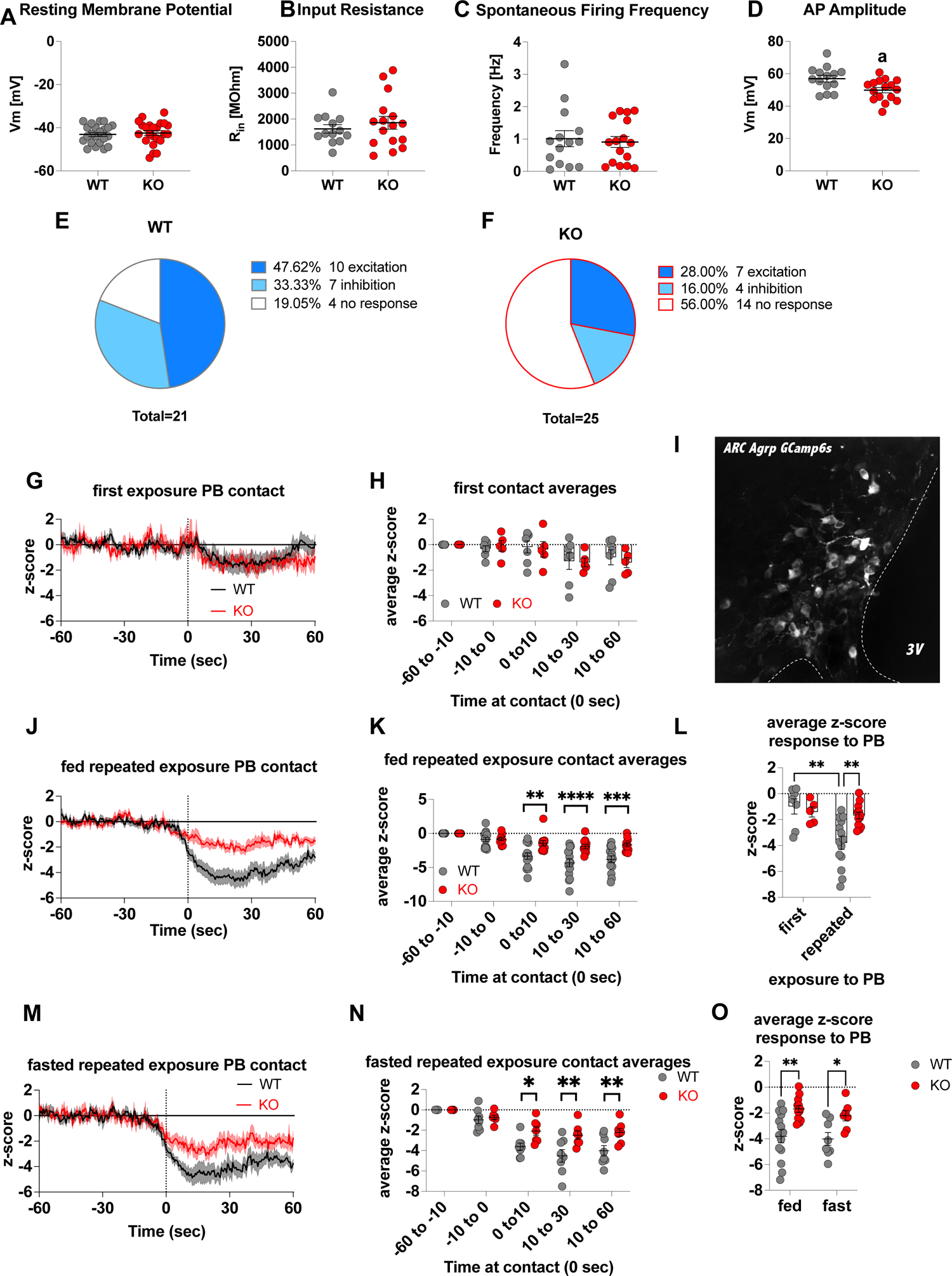
Deletion of Crat in AgRP neurons is a model of impaired metabolic responsiveness. Intrinsic electrophysiological properties were little affected in KO versus WT mice including: resting membrane potential (A), input resistance (B), spontaneous action potential firing frequency (C), although action potential amplitude was significantly reduced in KO AgRP neurons (D). Glucose responsive profiles of AgRP neurons from WT and KO mice characterised by their response to increased extracellular glucose concentration from 2mM to 5mM (E-F). In vivo fiber photometry analysis of AgRP neuronal activity in WT and KO mice (G-O). First exposure to a peanut butter chip (PB) (G, n=7 WT; n= 5 KO) and time binned summary data (H). GCaMP6s expression in AgRP neurons (I). AgRP neural activity in ad libitum fed mice in response to repeated PB exposure (J; n=7 WT; n= 5 KO) and time binned summary data (K-L, n=14 WT, n=12 KO). AgRP neural activity in fasted mice in response to repeated PB exposure (M; n=8 WT; n= 7 KO) and time binned summary data (N), n=8 WT, n=7 KO). Time binned summary data comparing the response to repeated PB exposure in ad libitum fed or fasted WT and KO mice (O). Data +/- SEM; two-way ANOVA with Tukey’s post hoc analysis (E, F, J, M, N) and unpaired students t-test (A-D); a, significant at p<0.05; ** p<0.01, *** p<0.001, **** p<0.0001. Dashed lines in I and J indicated the moment of contact with a PB contact.

**Figure 2:**
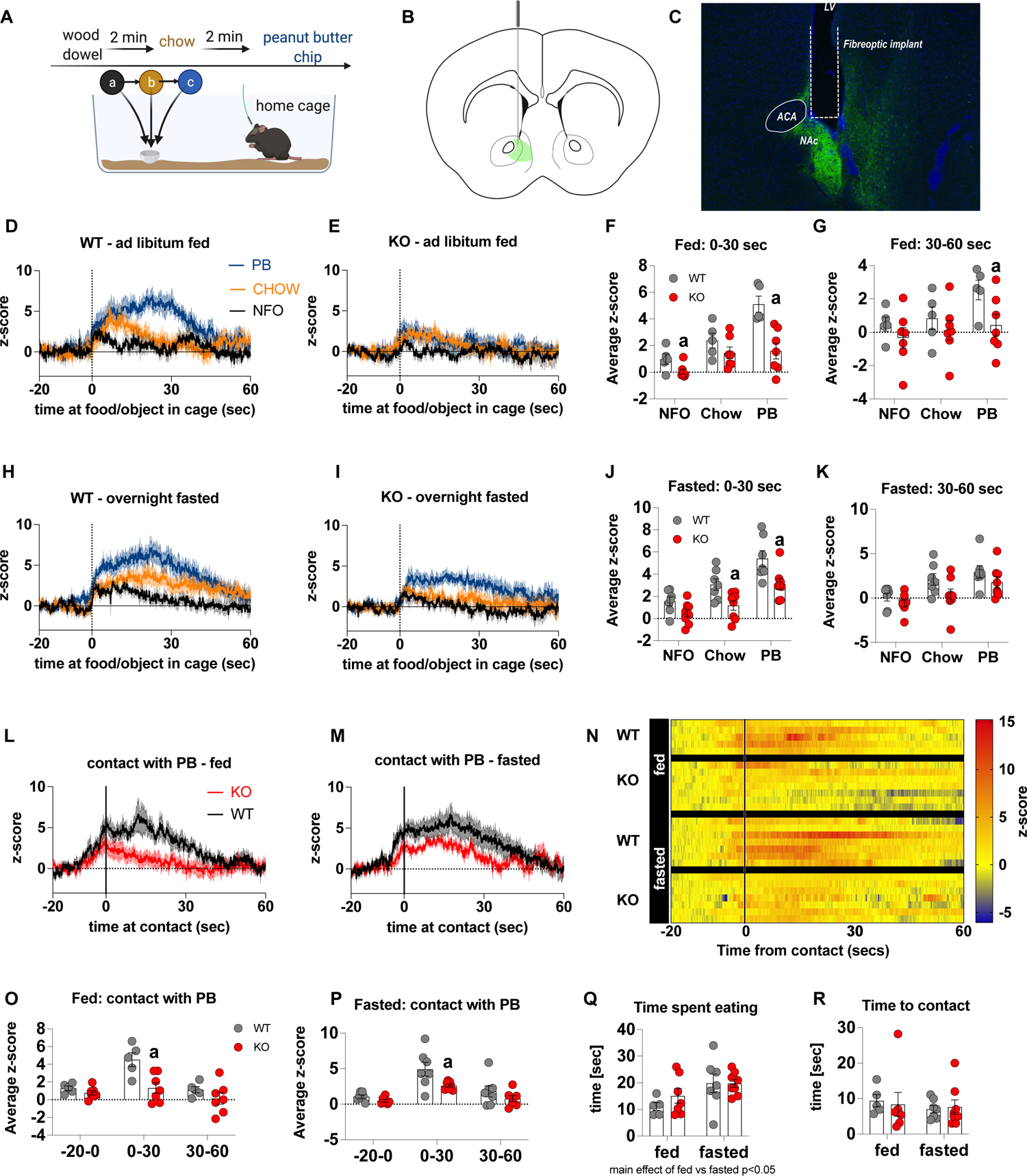
Impaired metabolic-sensing in AgRP neurons affects dopamine release in the nucleus accumbens. Experimental design to examine dopamine signalling in nucleus accumbens (A). Female (3 WT, 4 KO) and male (4 WT, 4 KO) mice, trained to receive peanut butter chips, were tethered to a fiber optic cable in their home cage. Following acclimation, dopamine responses in the nucleus accumbens to a non-food object (NFO; wood dowel, a, black), standard chow (b, tangerine), and peanut butter chip (PB, c, blue) were measured. Schematic of GRAB-DA (AAV-hSyn-DA4.3) injection site in the nucleus accumbens (B) and fiber placement (C). Average z-scored dopamine release traces aligned (time = 0) to object dropping into cage of WT and KO fed mice (D, E) or WT and KO fasted (H, I) mice. The average z-score of WT and KO ad libitum fed mice from 0-30 secs (F) and 30-60 secs (G) in response to a non-food object (NFO), standard chow, or peanut butter chip (PB) dropped into cage. The average z-score of WT and KO fasted mice from 0-30 secs (J) and 30-60 secs (K) in response to a non-food object (NFO), standard chow, or peanut butter chip (PB) dropped into cage. Average z-score dopamine release traces aligned to first contact with peanut butter chip (PB) from fed (L) and fasted (M) WT and KO mice, with individual traces shown in a heatmap (N). The average z-score of WT and KO ad libitum fed mice from −20-0, 0-30 and 30-60 secs (O), relative to first contact. The average z-score of WT and KO fasted mice from −20-0, 0-30 and 30-60 secs (P), relative to first contact. Time spent eating during the recording session (Q) and time passed between drop and contact with peanut butter (P). Data +/- SEM, two-way ANOVA with Tukey’s post hoc analysis (Q – main effect of fasting, p<0.05; R) and unpaired students t-test [F, G, J, K, O, P]; a – significant compared to WT, p<0.05. Dashed lines at time = 0 (D, E, H, I) represent the time at which food or an object was dropped into the cage. A continuous line at time = 0 (L, M) represents the time at which mice contact PB chip.

### Metabolic-sensing in AgRP neurons affects mesolimbic dopamine pathways

Previous studies show that chemogenetic activation of AgRP neurons modulates dopamine release in the NAc to food (*13*). To assess whether impaired metabolic-sensing in AgRP neurons affected dopamine release in the NAc, we used *in vivo* fiber photometry with the recently developed GRAB-DA sensor (*27*). Acute dopamine release in the NAc was measured in response to a non-food object (wood dowel), chow food and a peanut butter (PB) chip with each presentation separated by 2 mins (presented in that order) (Fig 2A). We observed a significant increase in dopamine release in response to chow or a PB chip presentation when compared to wood dowel in both fed and fasted WT mice 0-30 secs after placing the food or object into the cage (Fig 2D, E, H, I). Strikingly, KO mice had significant lower NAc dopamine release between 0-30 secs, as assessed by average z-score, after a non-food object or PB chip was placed in the cage compared to WT fed mice (Fig 2F). Average z-score was also significantly reduced in fasted KO mice in response to chow and PB chip (0-30 secs) when compared to WT fasted mice (Fig 2J). At 30-60 secs after food or an object was placed in the cage, the average z-score was only significantly lower in KO in response to PB in the fed mice (Fig 3G, K), suggesting the majority of the differential effects occur within the first 30 secs after presentation.

**Figure 3:**
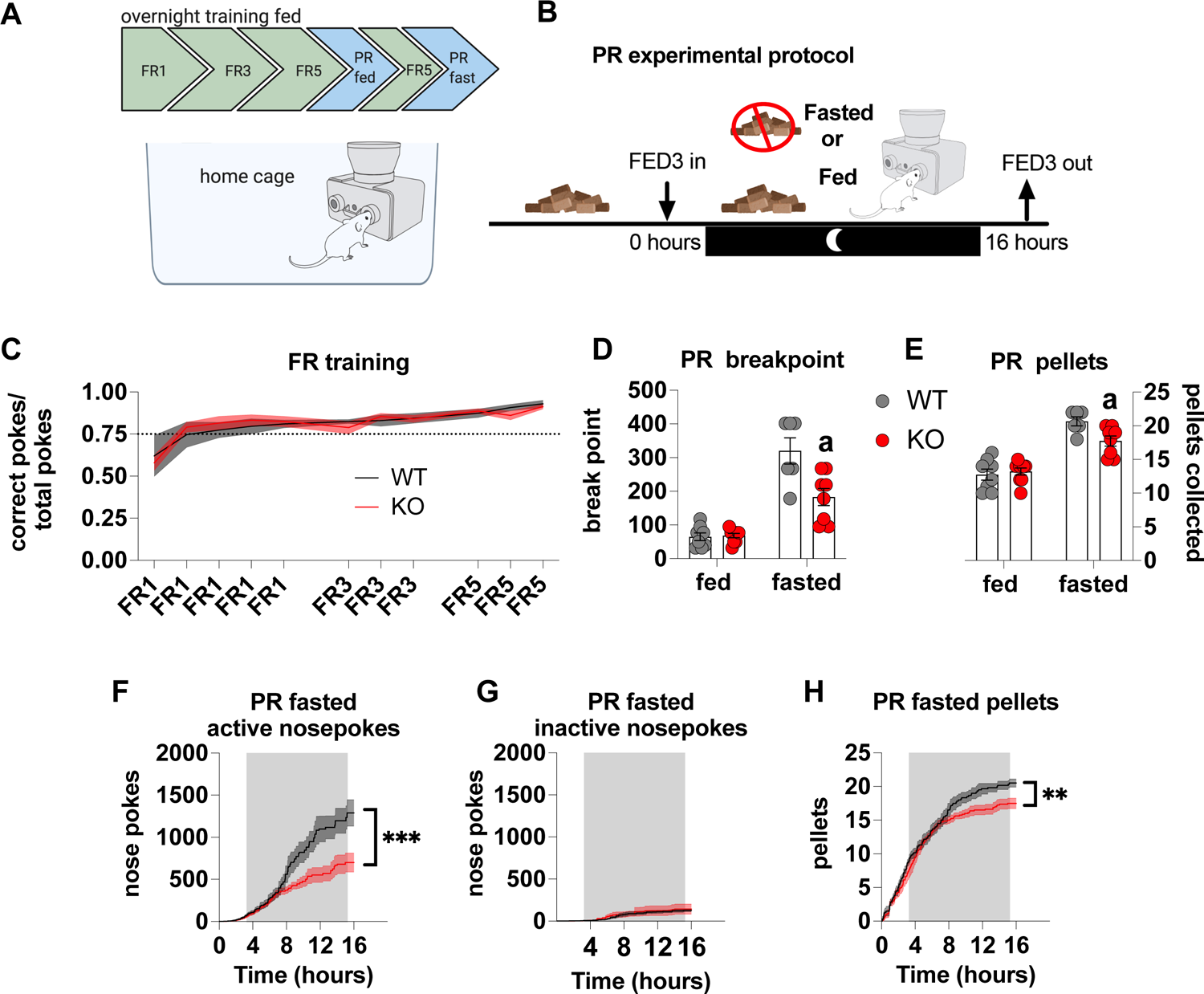
Impaired metabolic sensing in AgRP neurons affects motivation for sucrose rewards during fasting. Mice (8 WT, 9 KO) were trained to nose poke for sucrose rewards using fixed ratio schedule (A - FR1, FR3, FR5) to reliably nose poke on average 75% correct for 3 consecutive nights (C), before undergoing PR schedules with or without chow accessible. For PR under fasting conditions, mice were not fasted prior to testing but progressively fasted from the beginning until the end of the PR session (B). This important distinction tests the motivation to the progressively increasing energy deficit rather than in response to an already higher energy deficit. Breakpoint from fed and fasted mice (D) and pellet retrieval at the end of PR sessions (E). The number of active nose pokes (F), inactive (G) and pellets collected during PR fasted sessions. Data +/- SEM, two-way ANOVA with Tukey’s post hoc analysis (C, D, E, F – main effect of genotype, G, H – main effect of genotype). a – significant compared to WT, p<0.05, ** p<0.01, *** p<0.001.

Furthermore, we also aligned the dopamine signal to first contact with the PB chip (Fig 3L, M) and showed that there was a significantly reduced dopamine response 0-30 sec after contact with PB in KO compared to WT mice in both fed and fasted states (Figure O, P). No differences in dopamine release were observed in the 20 seconds prior to first contact or from 30-60 seconds after contact (Figure O, P). Taken together, these results highlight that impaired metabolic-sensing in AgRP neurons affects acute dopamine release in the NAc in both fed and fasted states in response to chow and a peanut butter chip. Time spent eating or time to first contact were not significantly different in WT and KO mice in fed or fasted state showing that these factors could not account for genotype differences in dopamine release in the NAc, although there was an expected main effect for fasting to increase overall time spent eating compared to fed mice, irrespective of genotype (Fig 3Q, R).

### Metabolic-sensing in AgRP neurons affects motivation and dopamine release during operant responding for a sucrose reward

The majority of operant conditioning protocols are conducted in stand-alone operant chambers and typically include a mild calorie restriction to facilitate learning the operant task. This exploits the fact that food restriction enhances appetitive drive, which is a result of elevated AgRP neuronal activity (*7, 12*). To avoid this potential confound we placed the open source FED3 (*30*) in home cages to facilitate learning in a low stress environment for long periods of time. As such, operant learning can be acquired quickly and easily in *ad libitum* fed mice without the need for caloric restriction – an important consideration for our experiments in which differential feeding responses to fasting, as reported previously (*25, 35*), may affect task acquisition and performance (Fig 3A). With this approach, there were no differences in operant responding during FR sessions (Fig 3C), indicating no differences in learning the operant task. In order to test motivation, progressive ratio sessions were performed overnight with (fed) and without access (fasted) to chow. Importantly, in fasted PR experiments, mice began fasting at the same time FED3s were placed in home cages (Fig 3B). This was designed to test the motivational response to an increasing energy deficit, which requires an awareness of homeostatic state over time, rather than in response to a prolonged fast. During PR fed experiments, WT and KO mice displayed similar breakpoint ratios (Fig 3D) and total number of pellets collected (Fig 3E), however during a fasted PR, KO mice showed a reduced breakpoint and total pellet consumption compared to WT mice (Fig 3D, E). Active (Fig 3F) but not inactive nose pokes (Fig 3G) during the fasted PR session were significantly lower in KO mice compared to WT, as well as pellets received during the progressive ratio session (Fig 3H). Intriguingly, active nose pokes and pellets collected only diverge around 6-8 hours after the beginning of fasting and the PR session, similar to when food intake diverges after fasting as previously described (*25, 35*). This timepoint likely reflects the stage at which impaired metabolic-sensing in AgRP neurons can no longer accurate report homeostatic state and energy need appropriately. Collectively, these results suggest that the appropriate detection of homeostatic state via metabolic-sensing in AgRP neurons affects motivation for a palatable food reward.

In order to establish that deficits in motivation were associated with impaired dopamine release in the NAc, we used FED3 as it allows for programmable TTL output to synchronize nose poking and pellet retrieval with measurements of dopamine release by GRAB-DA photometry (Fig 4A). Dopamine release was measured during a progressive ratio allowing alignment of dopamine release to rewarded or non-rewarded nose pokes. In fed and fasted WT mice, a rewarded nose poke significantly increased NAc dopamine release, as assessed by average z-score 0-60 secs after a poke event compared to 5 seconds prior to poke (Fig 4D, H, F, G). However, a rewarded nose poke in fed or fasted KO mice did not significantly increase NAc dopamine release within 60 seconds of the nose poke compared to 5 seconds prior to poke (Fig 4E, I, F, G). In addition, NAc dopamine release was significantly reduced during the 60 second period after a rewarded nose poke in KO compared to WT mice in both fed and fasted states (Fig 4F, G, J). The time between nose pokes was lower in fasted mice compared to fed mice (main effect; Fig 4K), but no genotype differences were observed.

**Figure 4.**
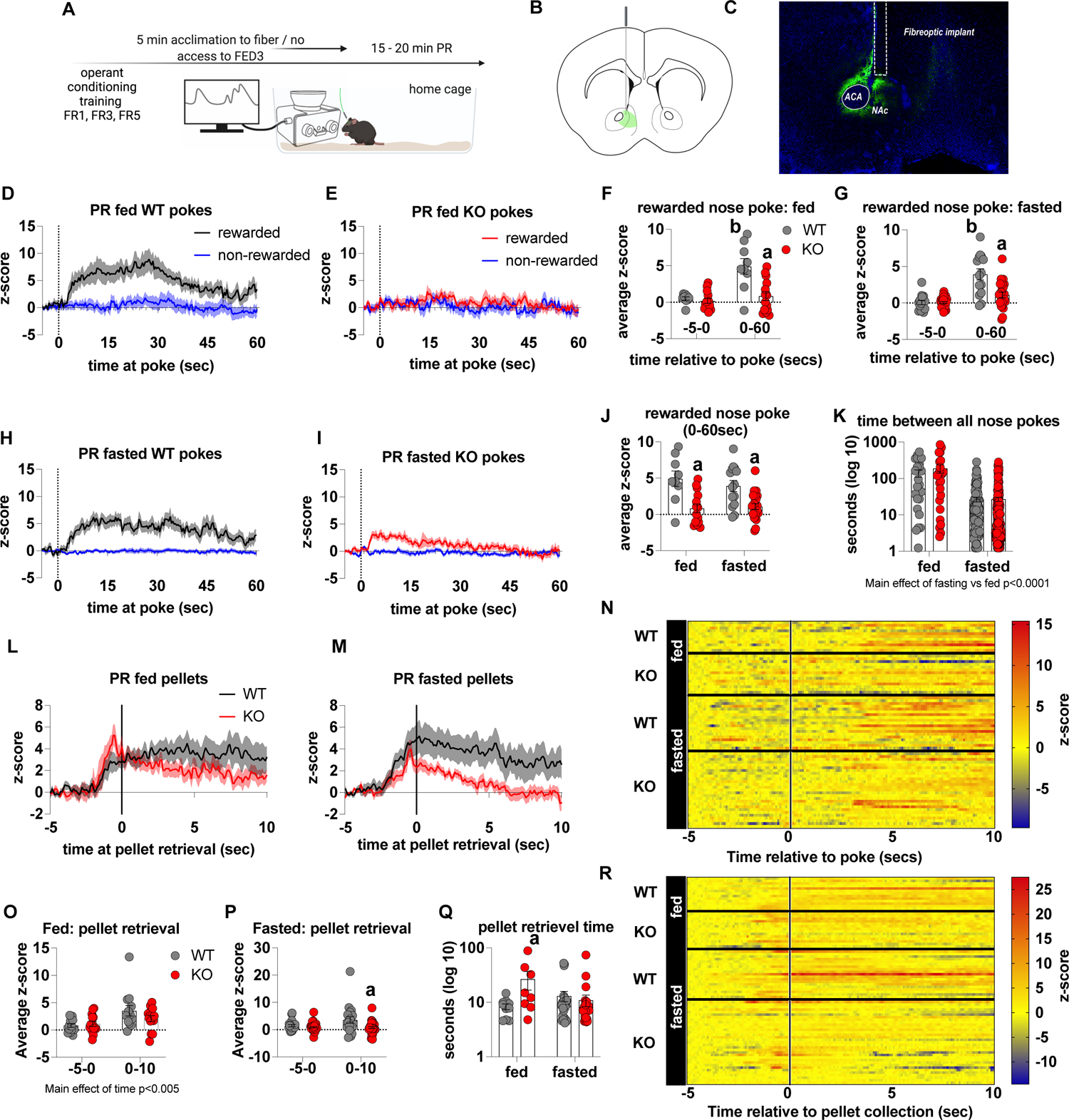
– Impaired metabolic sensing in AgRP neurons reduces dopamine release in the nucleus accumbens during a progressive ratio session. Experimental design (A) - mice with fiberoptic implant in the nucleus accumbens were trained to nose poke for a sucrose pellet prior to experimental testing. During testing, mice were tethered to a fiberoptic cable in their home cage and after 5 min acclimation, mice gained access to FED3 on PR schedule. Schematic of GRAB-DA (AAV-hSyn-DA4.3) injection site in the nucleus accumbens (B) and fiber placement (C). Combined average dopamine release response aligned (time = 0) to rewarded and non-rewarded correct nose pokes during a PR session from WT (D; n=4) and KO (E; n=4) ad libitum fed mice and WT (H; n=7) or KO (I, n=6) after an overnight fast. The average z-score of rewarded nose pokes from WT and KO ad libitum fed mice at −5-0 secs and 0-60 secs (F) or from WT and KO fasted mice at −5-0 secs and 0-60 secs (G). The average z-score of rewarded nose pokes at 0-60 secs in WT and KO mice during ad libitum or fasted conditions (J). The time between rewarded and non-rewarded nose pokes during the recording PR session in ad libitum fed or fasted WT and KO mice (K). Heat maps of individual responses from experimental mice aligned to nose poke (N). Average z-score dopamine release traces aligned to pellet retrieval (time =0) from ad libitum (L) or fasted (M) WT and KO mice. The average z-score at −5-0 secs and 0-10 secs after pellet retrieval in ad libitum fed (O) or fasted (P) WT and KO mice with pellet retrieval time for fed and fasted WT and KO mice shown in Q. Heatmaps of individual responses from experimental mice aligned to pellet retrieval (R). Data +/- SEM, two-way ANOVA with Tukey’s post hoc analysis (F, G, J, K – main effect of fasting, O - main effect of time, P, Q). a – significant compared to WT, p<0.05, b – significant compared to WT −5-0 secs, p<0.05. Dashed lines at time = 0 (D, E, H, I) represent the time at which a nose poke was made. A continuous solid line at time = 0 (L, M) represents the time at which mice collected the pellet from the pellet dispenser.

We also analysed dopamine release aligned to pellet retrieval (Fig 4L, M, O, P, Q, R) and observed no genotype differences in the fed mice (Fig 4L, O), although there was a main effect for increased NAc dopamine release 0-10 seconds after pellet retrieval compared to 5 seconds immediately prior to pellet retrieval. In the fasted state, dopamine release was significantly lower 0-10 seconds after pellet retrieval in KO mice compared to WT mice (Fig 4P). Interestingly, pellet retrieval was significantly longer in fed KO mice compared to fed WT mice (Fig 4Q). These results are consistent with progressive ratio data, in which breakpoint, active nose pokes and pellets collected are all significantly lower in fasted, but not fed, KO mice compared to WT mice. Thus, reduced NAc dopamine release in response to pellet retrieval presumably underpins reduced motivation for a palatable food reward in fasted KO mice.

Previous studies have suggested that neural encoding of non-caloric and caloric solutions are differentially processed in the NAc and dorsal striatum respectively (*36*). Next, we explored dopamine release in the dorsal striatum of WT and KO mice under fed and fasted conditions. We used the same protocol to measure acute dopamine release in the dorsal striatum in response to a non-food object (wood dowel), chow food and a PB chip with each presentation separated by 2 mins (Fig 5A). Although we observed a main effect for a PB chip to acutely increase dopamine release in the dorsal striatum 0-30 seconds and 30-60 seconds after food or object presentation, there were no significant genotype differences in either the fed or fasted state (Fig 5D-K). When data was aligned to contact with PB, we observed no genotype differences in contact with PB in fed or fasted mice at any time point (−20-0, 0-30, 30-60 secs; Fig 5L, M, S). However, we observed a main effect of time to increase dopamine release at 0-30 secs in fed and fasted mice (Fig 5O, P) independent from genotype showing that PB consumption elicits an acute increase in dopamine release in the dorsal striatum. Similar to data collected from the NAc experiments, no differences in time to contact with a PB chip or time spent eating a PB chip (5Q, R) were observed.

**Figure 5.**
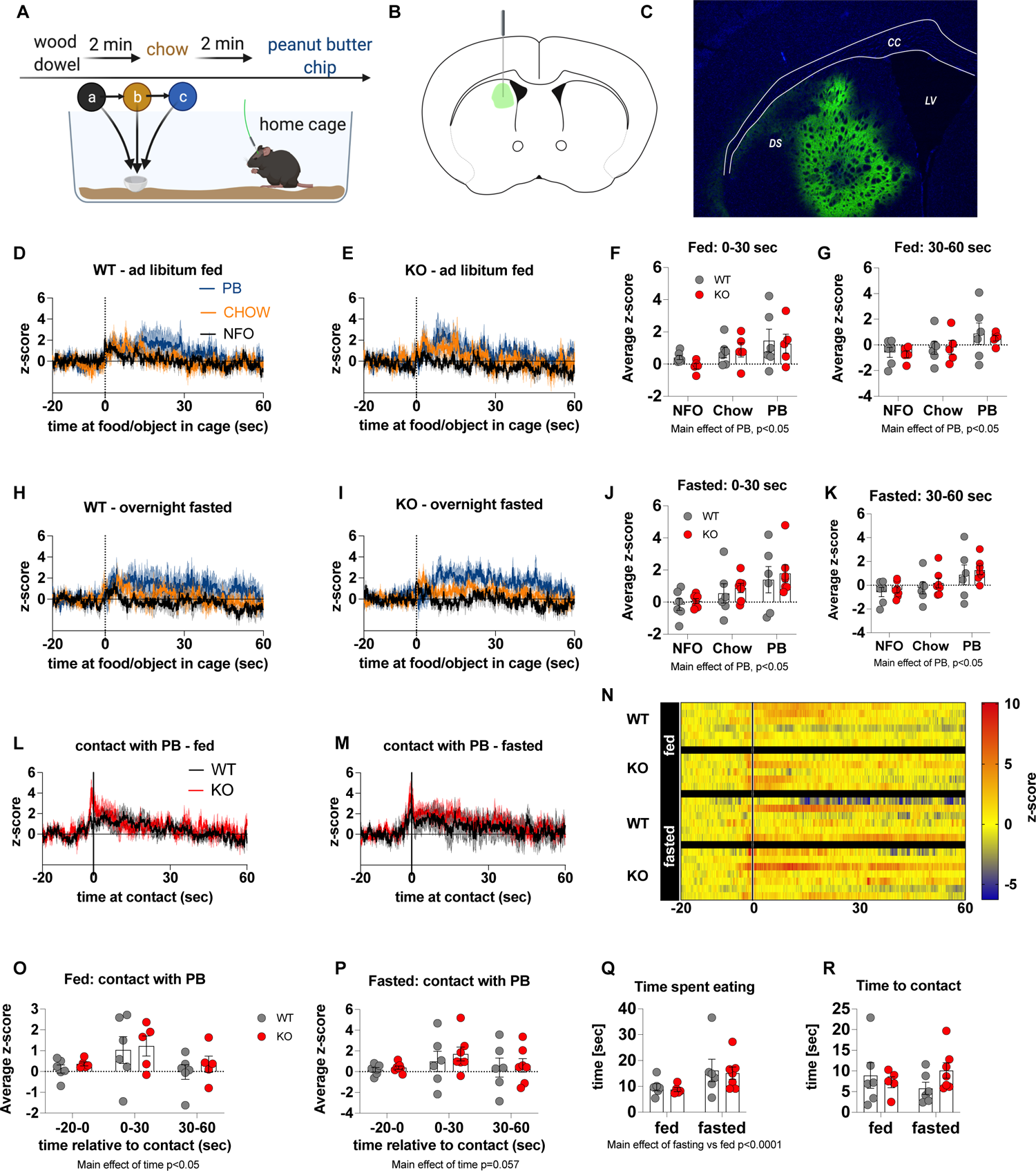
The effect of impaired metabolic-sensing in AgRP neurons on dopamine release in the dorsal striatum. Experimental design to examine dopamine signaling in dorsal striatum (A): Female (2 WT, 3 KO) and male (4 WT, 4 KO) mice, trained to receive PB chips, were tethered to a fiber optic cable in their home cage. Following acclimation, dopamine responses in the dorsal striatum to a non-food object (NFO; wood dowel, a, black), standard chow (b, tangerine), and peanut butter chip (PB, c, blue) were measured. Schematic of GRAB-DA (AAV-hSyn-DA4.3) injection site in the dorsal striatum (B) and fiber placement (C). Average z-scored dopamine release traces aligned (time = 0) to object dropping into cage of WT and KO fed mice (D, E) or WT and KO fasted (H, I) mice. The average z-score of WT and KO ad libitum fed mice from 0-30 secs (F) and 30-60 secs (G) in response to a non-food object (NFO), standard chow, or peanut butter chip (PB) dropped into cage. The average z-score of WT and KO fasted mice from 0-30 secs (J) and 30-60 secs (K) in response to a non-food object (NFO), standard chow, or peanut butter chip (PB) dropped into cage. Average z-score dopamine release traces aligned to first contact with peanut butter chip (PB) from fed (L) and fasted (M) WT and KO mice, with individual traces shown in a heatmap (N). The average z-score of WT and KO ad libitum fed (O) and fasted mice from −20-0, 0-30 and 30-60 secs (P), relative to first contact. Time spent eating during the recording session (Q) and time passed between drop and contact with peanut butter (R). Dashed lines at time = 0 (D, E, H, I) represent the time at which food or an object was dropped into the cage. A continuous line at time = 0 (L, M) represents the time at which mice contact PB chip. Data +/- SEM, two-way ANOVA with Tukey’s post hoc analysis (F, G, J, K – main effects of PB, p<0.05; O – main effect of time, p<0.05; P – main effect of time p=0.057; Q – main effect of fasting vs fed p<0.05).

We also examined dopamine release in the dorsal striatum during a progressive ratio schedule (Fig 6A). Aligning dorsal striatal dopamine to nose pokes showed a significant increase in dopamine release 0-10 seconds after a rewarded nose poke in both fed and fasted mice, independent from genotype (main effect of time; p<0.05; (Fig 6D-I). When aligned to pellet retrieval, dorsal striatal dopamine peaked immediately prior to retrieval (Fig 6J-K) with no significant genotype differences in average z-score prior to or after pellet retrieval (Fig 6L, M). Nevertheless, there was a significant main effect for reduced dopamine release after pellet retrieval compared with prior to pellet retrieval in both fed and fasted mice, independent from genotype (main effect of time; p<0.05; Fig 7L, M). Our data collected from the dorsal striatum suggest that impaired metabolic-sensing in AgRP neurons does not affect acute dopamine release after food/object presentation (0-60 seconds) or during a PR (0-10 seconds). However, our photometry approach cannot assess longer term changes, which may be an important factor since a study using human subjects showed that there are two temporally and spatially distinct dopamine responses to ingesting a milkshake (*37*). An immediate orosensory response occurs in response to taste and a second delayed post-ingestive response occurs in response to calories, which is localised to the dorsal striatum. To investigate this, we employed a positron electron tomography (PET) method using radiolabelled 18F-fDOPA (Fig 6N). First, we measured dynamic basal dopamine uptake in ventral and dorsal striatum in fasted mice without reward presentation and observed no differences in fDOPA uptake (Supp Fig 3A, B) suggesting impaired metabolic-sensing in AgRP neurons does not affect baseline uptake parameters. However, in response to PB chip consumption, we detected an increase in fDOPA accumulation 30 minutes after starting in the dorsal striatum of WT but not in KO mice (Fig 6P) and no differences in the ventral striatum (Fig 6O). These studies suggest that impaired metabolic-sensing in AgRP neurons restricts a post-ingestive dopamine response, ∼30-35 mins after consumption of a PB chip consumption, in the dorsal striatum.

**Figure 6.**
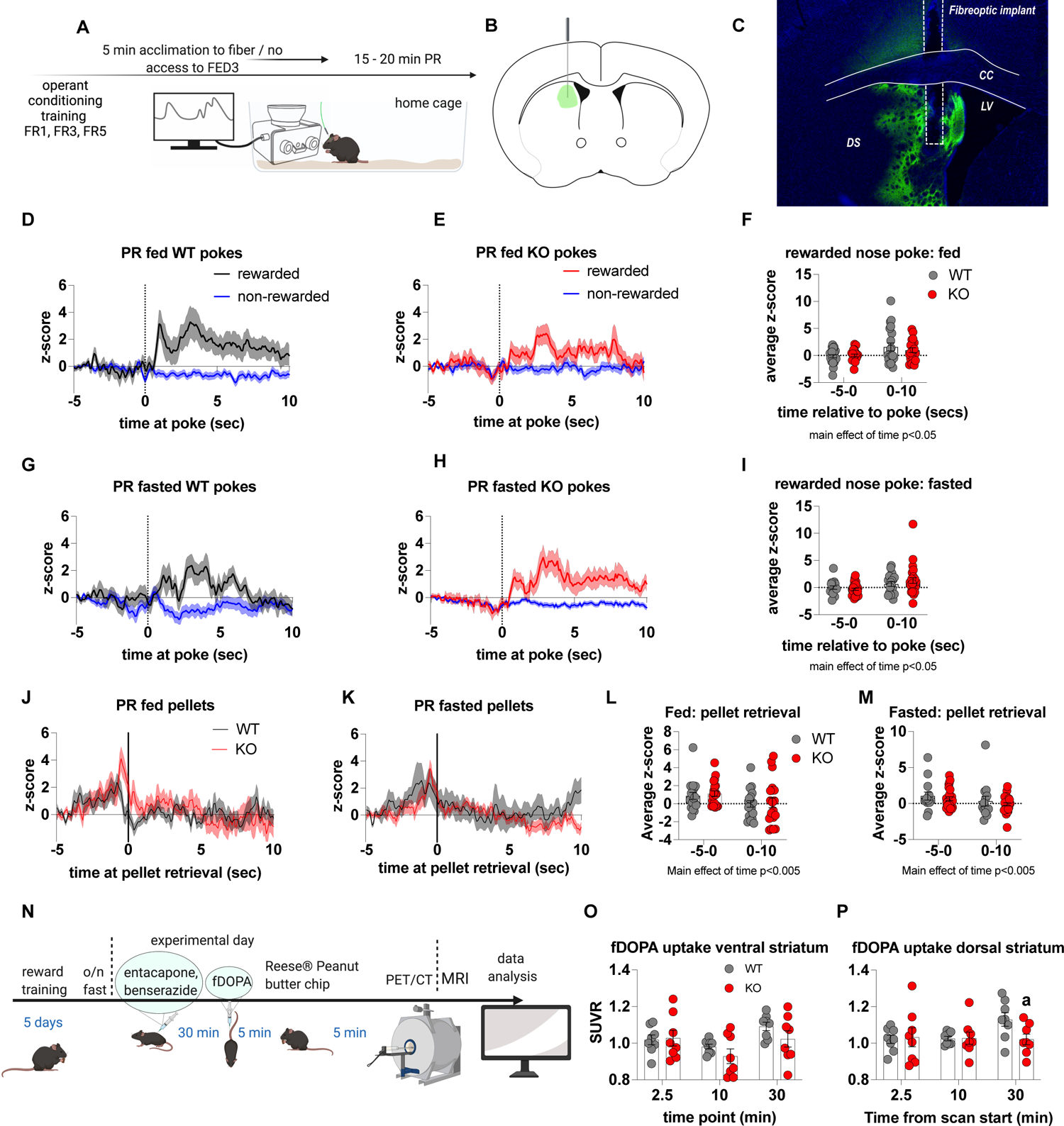
Impaired metabolic-sensing in AgRP neurons does not affect dopamine release in the dorsal striatum during a progressive ratio session. Experimental design (A) - mice with a fiberoptic implant in the dorsal striatum were trained to nose poke for a sucrose pellet prior to experimental testing. During testing, mice were tethered to a fiber optic cable in their home cage and after 5 min acclimation, mice gained access to FED3 on PR schedule. Schematic of GRAB-DA (AAV-hSyn-DA4.3) injection site in the dorsal striatum (B) and fiber placement (C). After 5 min acclimation mice gained access to FED3 on PR schedule. Combined average dopamine response of rewarded and non-rewarded correct nose pokes of 4 WT (D) and 4 KO (E) in fed state and 4 WT (G) and 4 KO (H) after overnight fast. The average z-score of rewarded nose pokes from WT and KO ad libitum fed mice at −5-0 secs and 0-10 secs (F) or from WT and KO fasted mice at −5-0 secs and 0-10 secs (I). Average z-score dopamine release traces aligned to pellet retrieval (time =0) from ad libitum (J) or fasted (K) WT and KO mice. The average z-score at −5-0 secs and 0-10 secs relative to pellet retrieval in ad libitum fed (L) or fasted (M) WT and KO mice. Experimental design for fDOPA PET scan (N): Mice were trained to receive Reese® peanut butter (PB) chips and then fasted overnight before the experimental day. On the experimental day, mice received ip injection of benserazide and entacapone to prevent peripheral breakdown of fDOPA (iv injection 30 min later). Mice were allowed to recover 5 min and then given 1 peanut butter chip, which they ate within 5 min. Then mice were anaesthetised and prepared for PET/CT scan and dopamine uptake in dorsal and ventral striatum measured. Baseline fDOPA uptake in ventral striatum (O) and dorsal striatum (P). Dashed lines at time = 0 (D, E, G, H) represent the time at which a nose poke was made. A continuous line at time = 0 (J, K) represents the time at which mice collected the pellet from the pellet dispenser. Data +/- SEM, two-way ANOVA with Tukey’s post hoc analysis (F, I, L, M – main effect of time, p<0.05). a – significant compared to WT, p<0.05.

## Discussion

In this study, we show that AgRP neurons require appropriate metabolic sensing of available calories in order to engage midbrain dopamine circuits and increase food motivation. We demonstrate that Crat deletion in AgRP neurons is a valid model of abnormal metabolic-sensing since it impairs glucose-sensing using *ex vivo* electrophysiological recordings and impairs *in vivo* AgRP GCaMP6 responses to repeated palatable food intake. An important role for Crat in metabolic sensing is supported by studies deleting Crat from myocytes, as this diminishes the switch from fatty acid metabolism to glucose metabolism after pyruvate administration (*38*).

There has been a shift in the understanding of AgRP neuronal function since it was discovered that the integration of sensory information suppresses AgRP neuronal activity prior to food consumption (*11, 18, 21*). Intriguingly, in our studies the first presentation of PB to WT and KO mice produces a similar reduction in AgRP neural activity suggesting normal integration of sensory and taste information, in the absence of post-ingestive interoceptive information. The subsequent fall in AgRP activity from WT mice after initial PB consumption shows that WT mice integrate additional interoceptive information based on post-ingestive feedback from PB consumption. However, KO mice do not show any greater suppression of AgRP neural activity with repeated PB exposure and consumption. Our results are supported by a number of studies illustrating that post-ingestive feedback from gut-derived signals is required to maintain the suppression of AgRP activity in response to consumption (*22–24*). In particular, Su et al demonstrated that the consumption of caloric gels, but not non caloric gels, is required for the sustained suppression of AgRP neuron activity, the magnitude of which is proportional to the calories obtained. In these studies the caloric value of a gel was learnt in a single trial (*22*), similar to the response seen in WT mice after first exposure to PB in our studies. The inability to sense caloric information in KO mice demonstrates a fundamental requirement for metabolic-sensing of AgRP neurons to link interoceptive caloric information to food presentation and consumption. Moreover, this unique model of impaired metabolic sensing in AgRP neurons caused by Crat deletion is not simply due to a dysfunctional AgRP neurons, since both food intake and neural activity in response to ghrelin and standard chow access were not different between genotypes. Thus we have generated a valid model of impaired metabolic sensing in AgRP neurons to test the hypothesis that metabolic-sensing of homeostatic state in AgRP neurons regulates motivational dopaminergic pathways.

By using GRAB-DA photometry to measure dopamine release dynamics (*27*) in the NAc and dorsal striatum, we assessed the impact of impaired metabolic sensing in AgRP neurons on dopamine release in response to PB. Although both WT and KO mice spent equal time consuming the peanut butter chip, KO had significantly attenuated acute dopamine release in the NAc in fed or fasted state. By contrast, we did not see any difference in acute dorsal striatum dopamine release between WT and KO mice. Thus, metabolic sensing in AgRP neurons is required to potentiate acute dopamine release in the NAc, extending previous observations that artificial activation of AgRP neurons increases dopamine release in response to food (*13*).

Importantly, impaired metabolic-sensing in AgRP neurons also attenuated progressive ratio operant responding for sucrose rewards during fasting with KO mice showing reduced breakpoint ratio, active nose pokes and pellets consumed. Although the motivation to obtain food is associated with chemo- or opto-genetic activation of AgRP neurons (*7, 12*), our studies show metabolic sensing of homeostatic state is required for translating low energy availability into a dopamine-driven motivational action to consume sucrose. We used a home-cage operant self-administration approach by placing FED3 in the home cage of mice. This provides advantages such as reducing handling stress, but most importantly, mice did not require food restriction to learn the action-outcome (nose poke-sucrose pellet) contingency and there were no differences in acquisition rates between WT and KO mice. This was important because differential responses to food restriction, as shown previously (*25, 39*), may have impaired subsequent responding during the progressive ratio tests.

Using FED3 coupled with GRAB-DA photometry, we measured dopamine release in the NAc or dorsal striatum during a progressive ratio session in response to rewarded and non-rewarded nose pokes. In the NAc, dopamine release increased prior to pellet retrieval and is consistent with FED3-motor activity acting as an auditory cue for pellet delivery, given the well-known role of cue-induced dopamine release in the NAc (*40, 41*). Intriguingly, cue-evoked dopamine release in the NAc reflects expected dopamine-mediated reinforcement rather than the actual magnitude of dopamine neuronal activation (*41*). There were no differences in average z-score prior to pellet retrieval, however average z-score was reduced after pellet retrieval in KO mice under fasted conditions. These observations show that metabolic-sensing in AgRP influences not the response to the reward predicting cue, but to the reward, which is driven by taste and caloric value, as previously described (*21, 24, 42*). The combination of reduced breakpoint responding and lower dopamine release in the NAc to pellet retrieval is consistent with the known role of NAc dopamine release to drive food seeking (*43*) and to assign an appropriate investment of effort to the available reward (*44–46*). In the dorsal striatum, dopamine ramping was observed prior to pellet retrieval without any genotype differences, which dropped to baseline soon after pellet retrieval consistent with previous reports (*47*).

With photometry, we were unable to detect any differences in dopamine release in the dorsal striatum yet studies in humans show palatable food elicits an immediate orosensory dopamine response and a delayed dorsal striatum dopamine response (*37*). Our photometry approach only examined a 60 second window after PB consumption or a 10 second window after pellet retrieval. Indeed after these short time periods dopamine release had already returned to baseline, making it impossible to utilise photometry to measure long-term changes in dopamine release. In order to address longer term changes in dopamine release we used 18F fDOPA in PET/CT studies and revealed reduced dopamine uptake 30 minutes after food reward in fasted KO mice compared to WT mice, with no difference in the ventral striatum. Thus we have identified an interesting separation in the temporal relationship of metabolic sensing in AgRP neurons and dopamine release in the NAc or dorsal striatum. Impaired metabolic sensing in AgRP neurons causes an acute attenuation of dopamine release in the NAc but not the dorsal striatum whereas it causes a longer-term attenuation in the dorsal striatum but not the ventral striatum, which encompasses the NAc. The exact mechanisms for these differences are unknown, although it is encouraging to see a similar temporal separation in an orosensory dopamine response and a delayed dorsal striatum dopamine response in humans (*37*). It is possible that the dorsal striatum response is delayed compared to the NAc response due to a requirement for metabolic processing in the gut or peripheral tissues. For example, disrupting peripheral glucose oxidation suppresses dorsal striatum dopamine efflux during sugar intake (*48*) and we have previously observed impaired glucose oxidation in response to refeeding after fasting in mice lacking Crat in AgRP neurons (*25, 35, 39*). Moreover, AgRP neurons respond to sensory cues within seconds yet require a slightly longer time frame (∼minutes) to response effectively to post-ingestive signals from the gut, such as nutrients and hormones released into the blood or via neural pathways (*22–24, 49*). Thus, we predict that impaired metabolic sensing in AgRP neurons affects both the immediate acute response to reward consumption, via dopamine signalling in the NAc, as well as slower response, which involves the integration of post-ingestive signals of calorie content in regions such as the dorsal striatum. In support of this, dopamine release in the dorsal striatum depends on calorie content, independent from taste, and is modulated by direct calorie infusion into the gut and gut-derived vagal afferents (*36, 50–52*).

Furthermore, we predict that both short term NAc dopamine release and longer term dorsal striatum dopamine release may be required to manifest a behavioural change in motivation, both of which require metabolic sensing in AgRP neurons. This comes from our observations that NAc dopamine release to a rewarded nose poke is lower in both fed and fasted KO mice, however reduced motivation for sucrose pellets in the PR session only manifests 6 hours into the fasted period. Indeed, we only observed changes in fDOPA accumulation in the dorsal striatum in mice fasted overnight.

An unanswered question remains, as to how AgRP neurons influence mesolimbic dopamine signaling. Given that very few, if any, AgRP terminals are found in the VTA (*14*), it seems unlikely that AgRP neurons directly affect dopaminergic neurons in the VTA, despite the increase in VTA dopamine activity in response to AgRP activation, which is presumably driven through indirect pathways (*15*). In addition, dopamine dynamics differentially regulate learning and motivational properties, with dopamine release in NAc underlying motivational properties and VTA firing properties underlying learning properties (*46*). Although the MC4R is found in the NAc (*53*), there is little evidence to suggest activation of AgRP neurons directly target neurons in the NAc. Activation of AgRP nerve terminals in the lateral hypothalamus (LH), but not the paraventricular hypothalamic nucleus, increased taste reactivity to sucrose (*42*), suggesting the LH may be an indirect route, through which AgRP neurons influence mesolimbic dopamine release. This is consistent with the known role of the LH in mediating appetitive motivation and food consumption (*54*). Finally, the suppression of AgRP neuronal activity with food presentation occurs prior to consumption (*11, 18, 21*) and this coincides with increased NAc and dorsal striatum dopamine release prior to pellet consumption. Whether or not these changes are functionally related remains to be determined but activation of AgRP neurons also drives dopamine release in the NAc (*13*) so a number of questions remain to be answered, as to how AgRP neurons facilitate dopamine release.

In summary, we show that metabolic sensing in AgRP neurons is required to transmit interoceptive metabolic information into dopamine release in the NAc and dorsal striatum, albeit over different time frames and to increase motivated behaviour for sucrose rewards. Since the motivation to eat depends not only on the palatability of food, but also metabolic state (*55, 56*), these studies identify a potential novel therapeutic strategy to control hunger-sensing AgRP neurons and prevent the overconsumption of palatable foods. Whether or not this plays an important role in obesity remains to be determined since obesity desensitizes AgRP neurons to food cues and metabolic feedback (*57*). However, the observation that fasting increases NAc dopamine release in response to high fat diet, when compared to chow diet (*15*), suggests that homeostatic circuits might play an important role in the pathogenesis of obesity. Future research is required to disentangle how homeostatic signals influence food reward processing and how this contributes to obesity.

## Acknowledgements

This study was supported by an NHMRC grant and fellowship to ZBA (1126724, 1154974). We would like to thank Myles Billard from TDT for his valuable technical assistance and support with setting up photometry and analysis. The authors acknowledge the facilities and scientific and technical assistance of the National Imaging Facility, a National Collaborative Research Infrastructure Strategy (NCRIS) capability, at Monash Biomedical Imaging, Monash University. We would like to thank Professor Alex Fornito for the use of fDOPA in PET studies. We acknowledge that Bio Render was used to produce elements incorporated in the figure and graphical abstract (Biorender.com).

## SUPPLEMENTARY FIGURE LEGENDS

Supplementary Figure 1 Sample traces showing examples of glucose excited (A) and glucose inhibited (B) neurons in response to an increase in glucose from 2 to 5mM glucose. Spontaneous excitatory (C) and inhibitory (D) post-synaptic potentials compared in WT and KO mice in response to an increase in glucose from 2mM to 5mM. Comparison of AgRP activity at first PB and repeated PB exposure in WT (E) and KO (F) mice (WT n=7 first exposure, n=14 repeated; KO n=5 first exposure, n=12 repeated exposure. Data +/- SEM

Supplementary Figure 2 Deletion of Crat from AgRP neurons does not affect ghrelin-induced food intake or AgRP activation. Cumulative food intake measured in BioDaq feeding cages over a 24-hour (A) and 4-hour time frame (B) in WT and KO mice treated with saline (WT n=5, KO n=7) and ghrelin (WT n=5, KO n=7). No genotypes difference in food intake were observed, however, ghrelin significantly increased food intake compared to saline (B, two way ANOVA, main effect of treatment). Experimental approach for neural recordings of AgRP activity in response to ghrelin and subsequent chow access (C). Note, mice were treated with IP ghrelin (1 mg/kg) for 90 minutes before chow access to prevent confounding interactions. IP ghrelin increased AgRP neural activity in both WT and KO (D) and although we observed a main effect for ghrelin (average z-score from 0-30 mins) to increase AgRP neural activity (E) relative to baseline (average z-score from −15-0 minutes), no genotype differences were observed (WT n=8; KO n=6; two way ANOVA). 90 minutes after IP ghrelin injection mice were given access to normal chow food, which was their primary daily food source. Food access resulted in an equal suppression of AgRP neural activity in WT and KO mice (F) and although we observed a main effect of food access to reduce AgRP activity, no genotypes differences were observed (WT n=8; KO n=6; two way ANOVA). No differences in food intake were observed in during the food access period (H). Light Dark box analysis and elevated plus maze analysis showed no genotype differences in time spent in the light zone or open arm, suggesting Crat deletion in AgRP neurons did not affect anxiety-like behaviour (WT n=14-15; KO n=8-10). Data +/- SEM, two-way ANOVA.

Supplementary Figure 3. Dynamic basal dopamine uptake in dorsal (A) and ventral striatum (B) in fasted mice without reward presentation and observed no differences in fDOPA uptake.

