## Supplementary figures and images for "Metabolic-sensing in AgRP neurons integrates homeostatic state with dopamine signalling in the striatum"

### Supp Fig 1

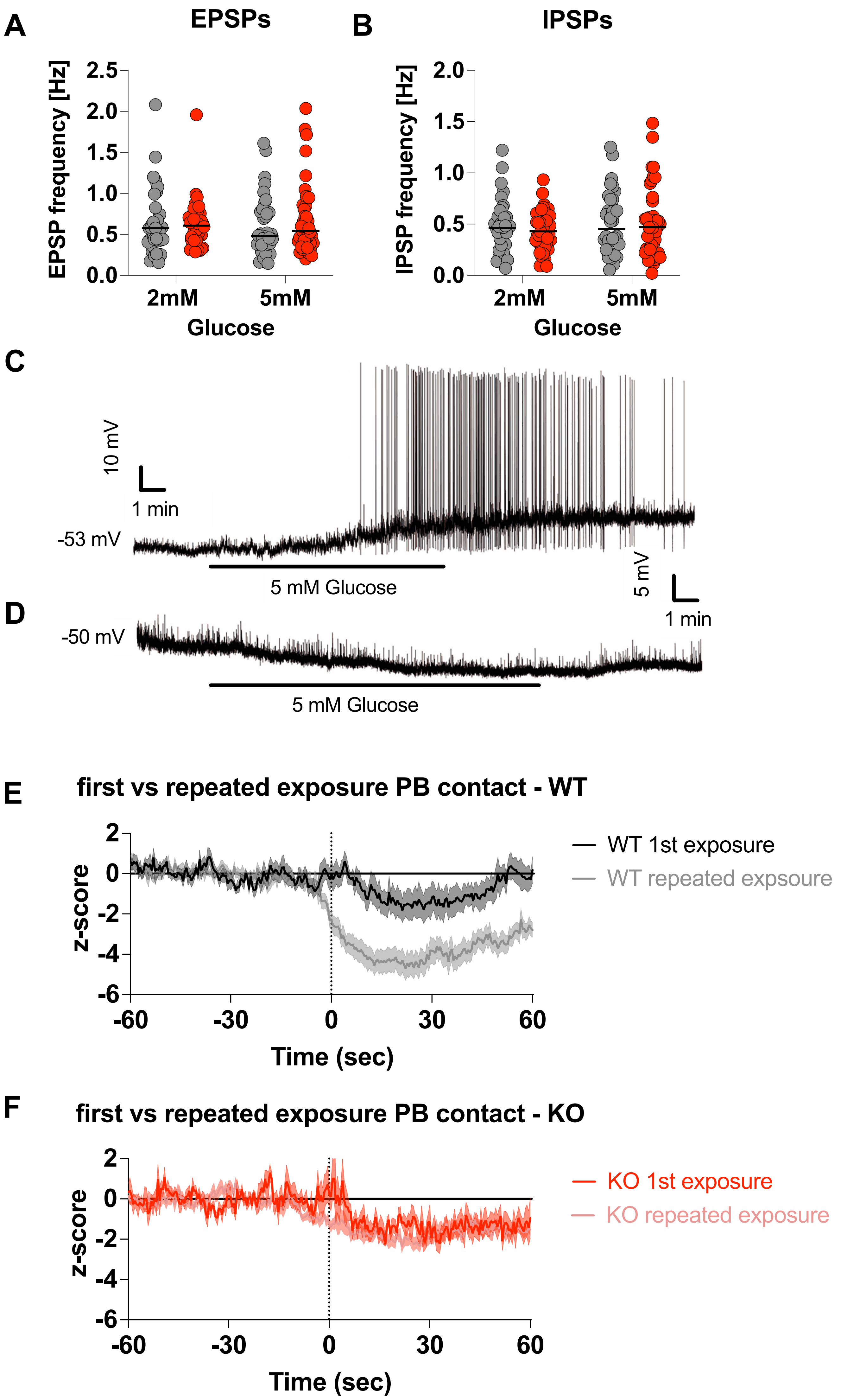

### Supp Fig 2

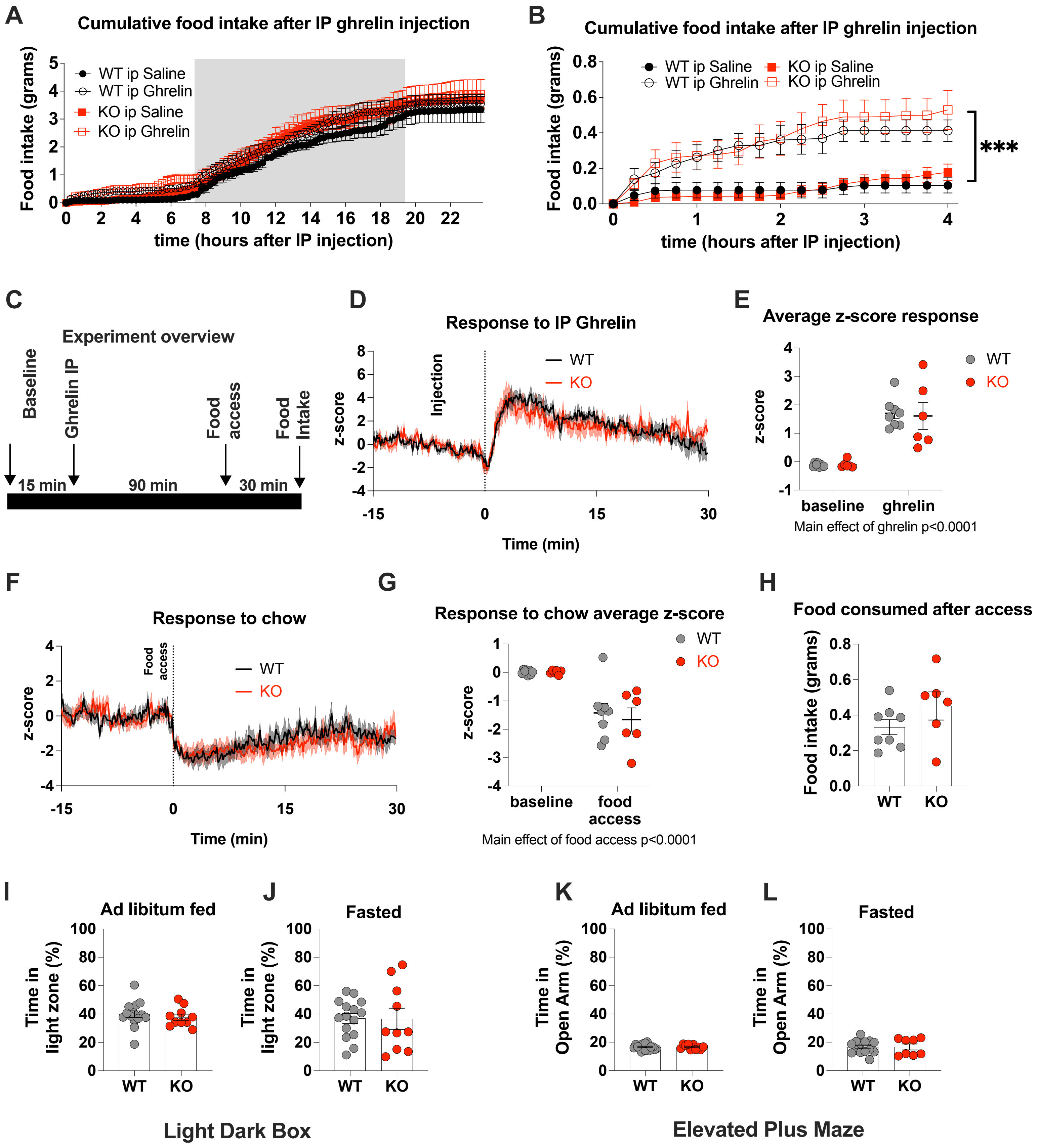

### Supp Fig 3

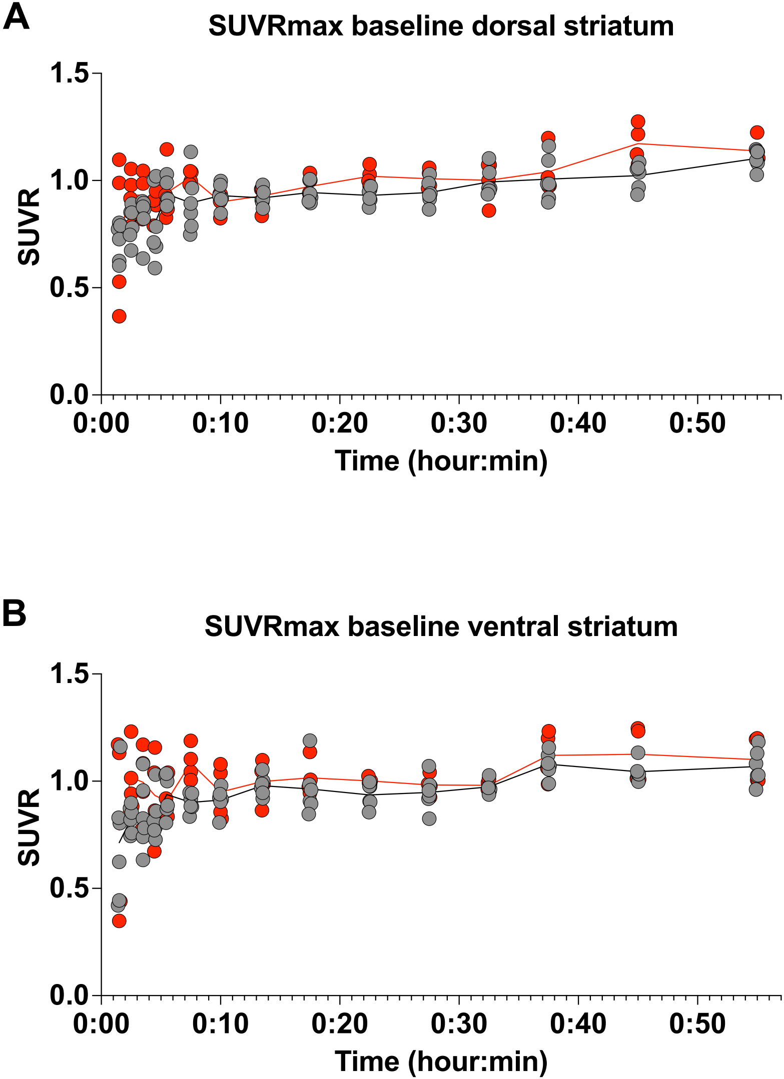
